# Signal-amplification for cell-free biosensors, an analog-to-digital converter

**DOI:** 10.1101/2023.04.14.536885

**Authors:** Rafael Augusto Lopes Franco, Gabriel Brenner, Vitória Fernanda Bertolazzi Zocca, Gabriela Barbosa de Paiva, Rayane Nunes Lima, Elibio Leopoldo Rech, Milca Rachel da Costa Ribeiro Lins, Danielle Biscaro Pedrolli

**Author notes:** Corresponding authors: Danielle B. Pedrolli, Milca Rachel da Costa Ribeiro Lins. Contributed equally as first author (Rafael Augusto Lopes Franco, Gabriel Brenner).

## Abstract

Toehold switches are biosensors useful for the detection of endogenous and environmental RNAs. They have been engineered to detect virus RNAs in cell-free gene expression reactions. Their inherent sequence programmability makes engineering a fast and predictable process. Despite improvements in the design, toehold switches suffer from leaky translation in the OFF state, which compromises the fold change and sensitivity of the biosensor. To address this, we constructed and tested signal amplification circuits for three toehold switches triggered by Dengue and Sars-CoV-2 RNAs and an artificial RNA. The serine integrase circuit efficiently contained leakage, boosted the expression fold-change from OFF to ON, and decreased the detection limit of the switches by three to four orders of magnitude. Ultimately, the integrase circuit converted the analog switches’ signals into digital-like output. The circuit is broadly useful for biosensors and eliminates the hard work of designing and testing multiple switches to find the best possible performer.

**GRAPHICAL ABSTRACT:** 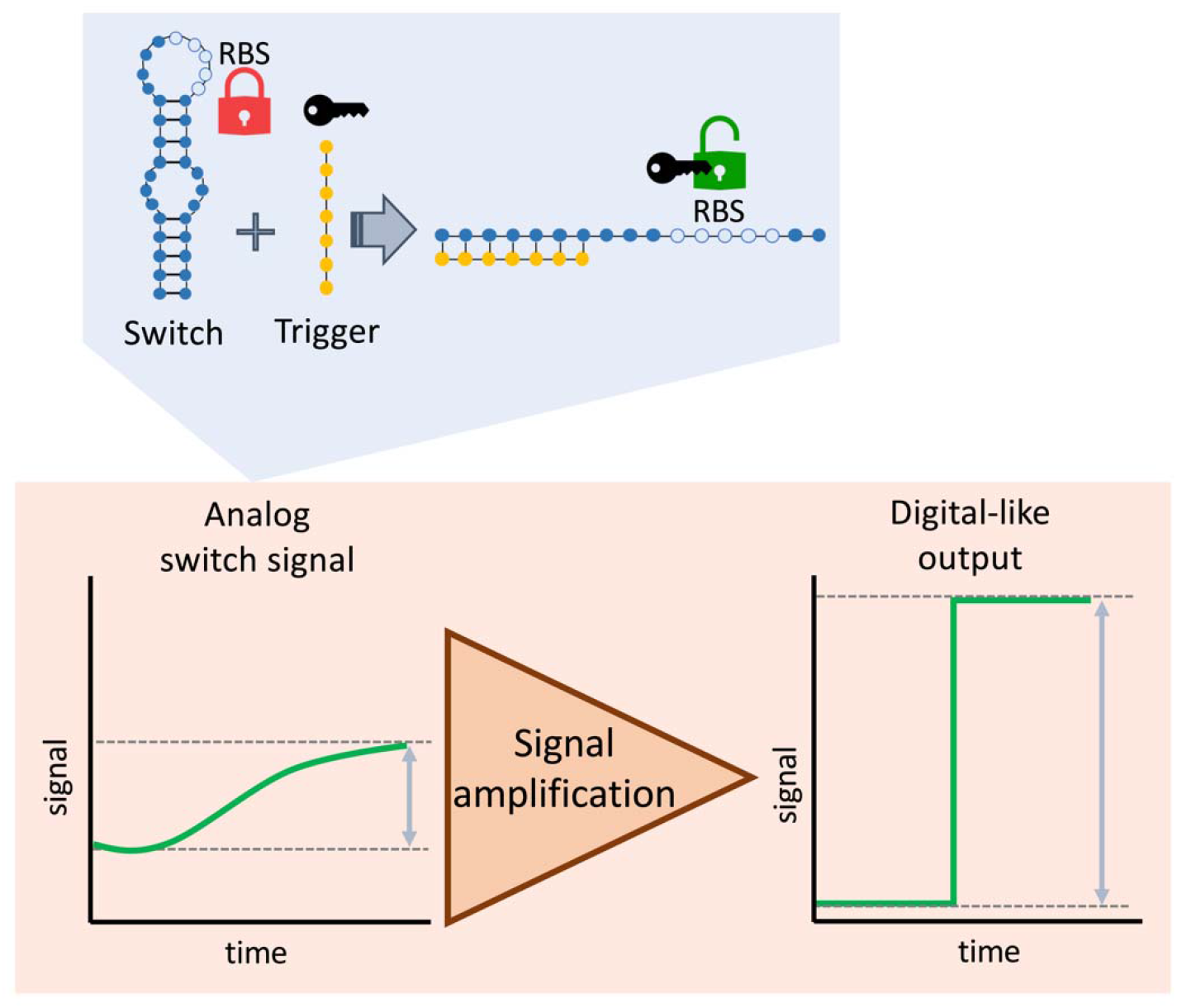

## 1. INTRODUCTION

Cell-free gene expression is a powerful tool for the study of gene expression in a controlled and customizable environment. It allows for the rapid synthesis of RNAs and proteins in a simple reaction with a small diversity of components, avoiding undesired contaminants and crosstalk. Additionally, it can be used to test and optimize the function of new sensors ^1,2^, circuits ^3,4^, and biosynthetic pathways ^5^. Finally, cell-free gene expression is an affordable and easy-to-implement system for biosensors. Protein sensors for water contaminants such as antibiotics, small molecules, and cations (including lead, arsenic, and mercury) have been successfully implemented using cell-free synthetic biology ^2,6^. For pathogens and human health biomarkers, RNA genetic switches have been used to detect specific target DNA or RNA sequences and trigger gene expression ^1,7^. Detection of RNA sequences has been particularly well studied using toehold switches rationally engineered to be triggered by RNAs of viruses such as Zika ^8^, Ebola ^1^, and Sars-CoV-2 ^9–11^, of pathogens in the human microbiome ^7^, and endogenous RNA of *E. coli* ^12^ and *Salmonella typhi* ^9^.

Toehold switches are synthetic regulatory RNAs able to control the mRNA translation. The toehold is a short sequence in the mRNA that is complementary to a target RNA molecule called the trigger. The toehold is attached to an RNA hairpin that sequesters the ribosome binding site (RBS) of the downstream gene in the mRNA. Consequently, the downstream gene cannot be translated. When the trigger RNA pairs with the toehold, it also invades the switch hairpin causing a strand displacement that releases the RBS and turns gene expression on. Toehold switches are valuable tools for engineering cells and cell-free systems, acting as RNA sensors and regulating gene expression ^1,12–14^.

Toehold switches are a powerful tool for designing new regulatory networks. RNA-based gene regulation presents advantages that make it particularly appealing for circuit engineering. Their inherent sequence programmability allows rational design and prediction of folding and molecular dynamics. Moreover, RNA regulators are compact and require only the transcription process ^12,15^. Compared to protein-based regulators, RNA regulators demand lower amounts of resources, impose lower production loads on the system, and can potentially propagate signals faster ^16^.

Despite many engineering efforts to improve the RNA switches, they still suffer from leaky translation in the OFF state, which compromises the fold change and sensitivity of the sensor ^8,9,13^. To circumvent the problem in cell-free reactions, an initial step of sample amplification is used such as NASBA ^7,8^ and RT-LAMP ^17^, which amplify clinically relevant concentrations of RNA to overcome the sensor’s detection limit. However, target trigger amplification is not feasible for *in vivo* applications such as whole-cell sensors and the detection of endogenous RNA species, limiting the detection to abundant target molecules only ^13^. One feasible solution for both cell-free and cellular systems is to use a cascaded circuit to amplify the transduced sensor signal and boost sensing performance. The solution has been successfully implemented in *E. coli* using transcriptional factors coupled with a protein sensor ^6^ and with a riboswitch ^18^.

To address the challenges related to low fold change and sensitivity of toehold switches, we designed and constructed three toehold switches triggered by Dengue RNA, Sars-CoV-2 RNA, and an artificial RNA. The switches were first tested in direct control of the reporter gene. In a first attempt to amplify the switch’s signal, we added a T7 RNA polymerase amplifier module to connect the switch to the reporter. The success in containing translation leakage came with a trade-off as a reduction in the maximum output production. Exchanging the T7 RNA polymerase for a serine integrase as an amplifier module changed the scenario completely. As a result, leakage was efficiently contained and the sensitivity was strongly boosted. The only trade-off detected is a delay in the response, which does not compromise the use of the sensor circuit.

## 2. RESULTS

### 2.1 Toehold Switches design and *in silico* analysis

Up to ten switch sequences were designed for each target RNA using NUPACK ^15^. Targets were unique 40 nt long sequences of RNA from Dengue type 1 and Sars-CoV-2 genomes and a non-specified RNA sequence. Only switch/trigger pairs showing 100% complex formation in the equilibrium at equimolar concentrations were selected ^15^. Next, the switches were analyzed for translation initiation rates in the OFF and ON states ^19–21^. Switches predicted to have translation rates up to 10^1^ in the OFF state and equal to or higher than 10^3^ in the on-state were selected and ranked according to their predicted dynamic range. Based on the *in-silico* analysis, the best-performing pair of toehold switch/target was constructed and tested (Supplementary tables 1 and 2).

### 2.2 Toehold Switch direct control of the output

Toehold switches have been designed and tested using simple devices where the switch directly controls the reporter gene. Different signals have been successfully used as outputs, such as GFP fluorescence ^1,12,14^, substrate color change catalyzed by beta-galactosidase ^1,8^, glucose release catalyzed by trehalase ^9^, and luminescence ^10^. We chose luminescence production catalyzed by an enhanced Luciferase (EPIC Luciferase) ^22^, which is highly sensitive although not visual. We tested our three switches’ ability to detect the RNA trigger and control the luciferase expression (Figure 1A). RNA triggers used were 40 nt-long synthetic RNAs mimicking genomic targets from the viruses Sars-CoV-2, Dengue, and Zika, and a synthetic RNA B. Switches’ specificity was accessed using relevant nonspecific target RNAs. All three switches were able to turn on gene expression when treated with the specific trigger in the micromolar range (Figure 1B-D). Switch B was activated by the specific trigger B at 7.2 µM, with a fold change of 2.9 after 3h (Figure 1B); however, lower concentrations failed in activating it. The specific Sars-CoV-2 trigger activated gene expression by 5.2-fold and 7.7-fold compared to the nonspecific triggers Sars-CoV-2 and Dengue, respectively (Figure 1C). The success achieved with triggers at 14.4 µM could not be reproduced with a lower concentration (Supplementary Figure 2A). The highest sensitivity was achieved with the Dengue switch, which was well able to discriminate between the specific trigger and the nonspecific Zika and SARS-CoV-2 triggers at 3.6 µM with a 2.8-fold change after 1h (Figure 1D). The nonspecific triggers tested are relevant to the Toehold switches’ activity as they may be found in samples used to detect the specific virus RNA as well. Although functional and specific, all three switches had low sensitivity as none of them could discriminate the specific trigger at concentrations lower than 3.6 µM (Figure 1 and Supplementary Figure 1), which is one order of magnitude higher than achieved for the detection of Zika and SARS-CoV-2 viruses RNA in other studies ^8,17^. At that point, we hypothesized that the expression leakage from our switches could account for the low sensitivities. To contain leakage and increase the output signal, we constructed gene circuits to transfer the switch signal to the output unit.

**Figure 1.**
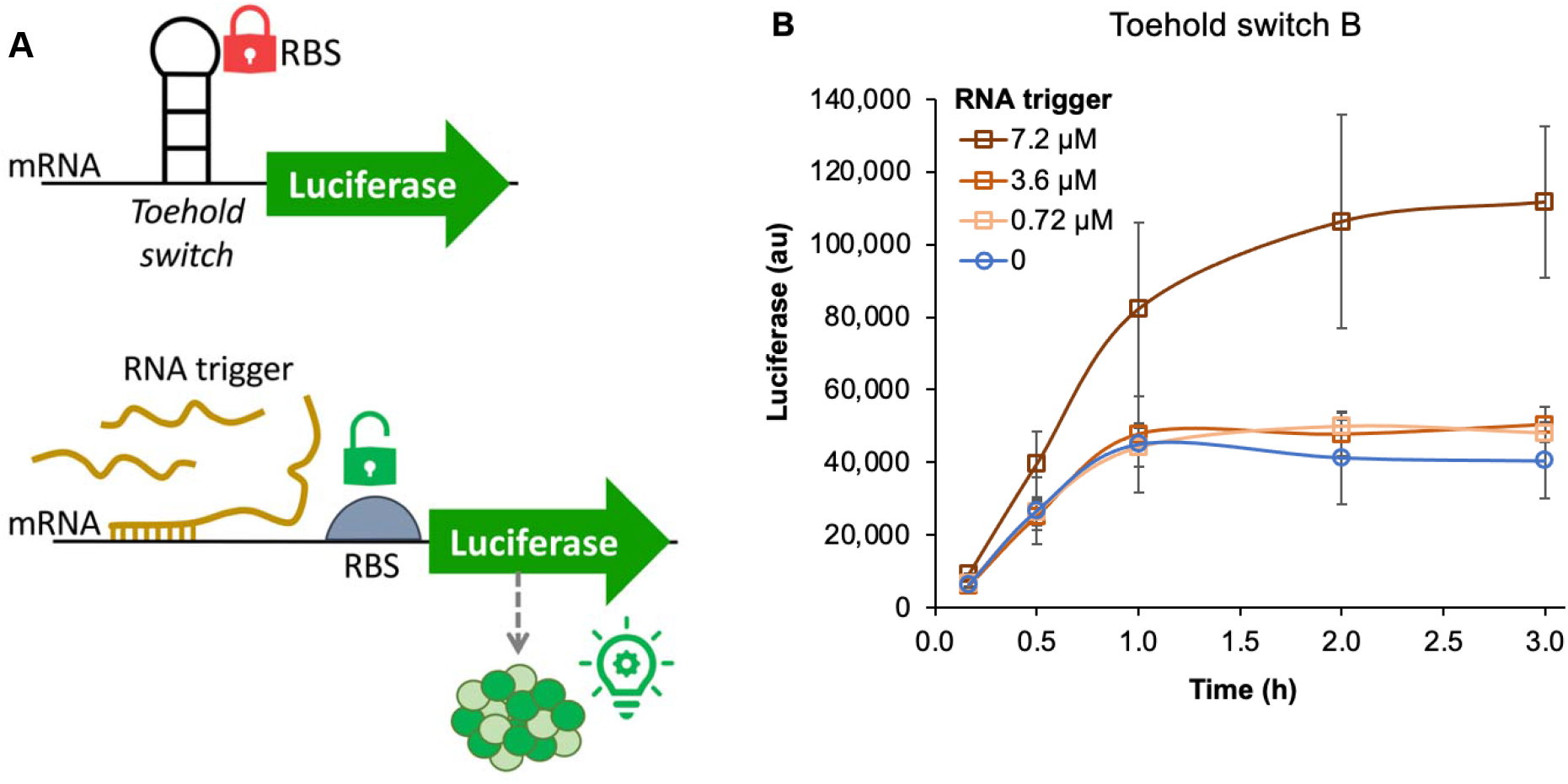
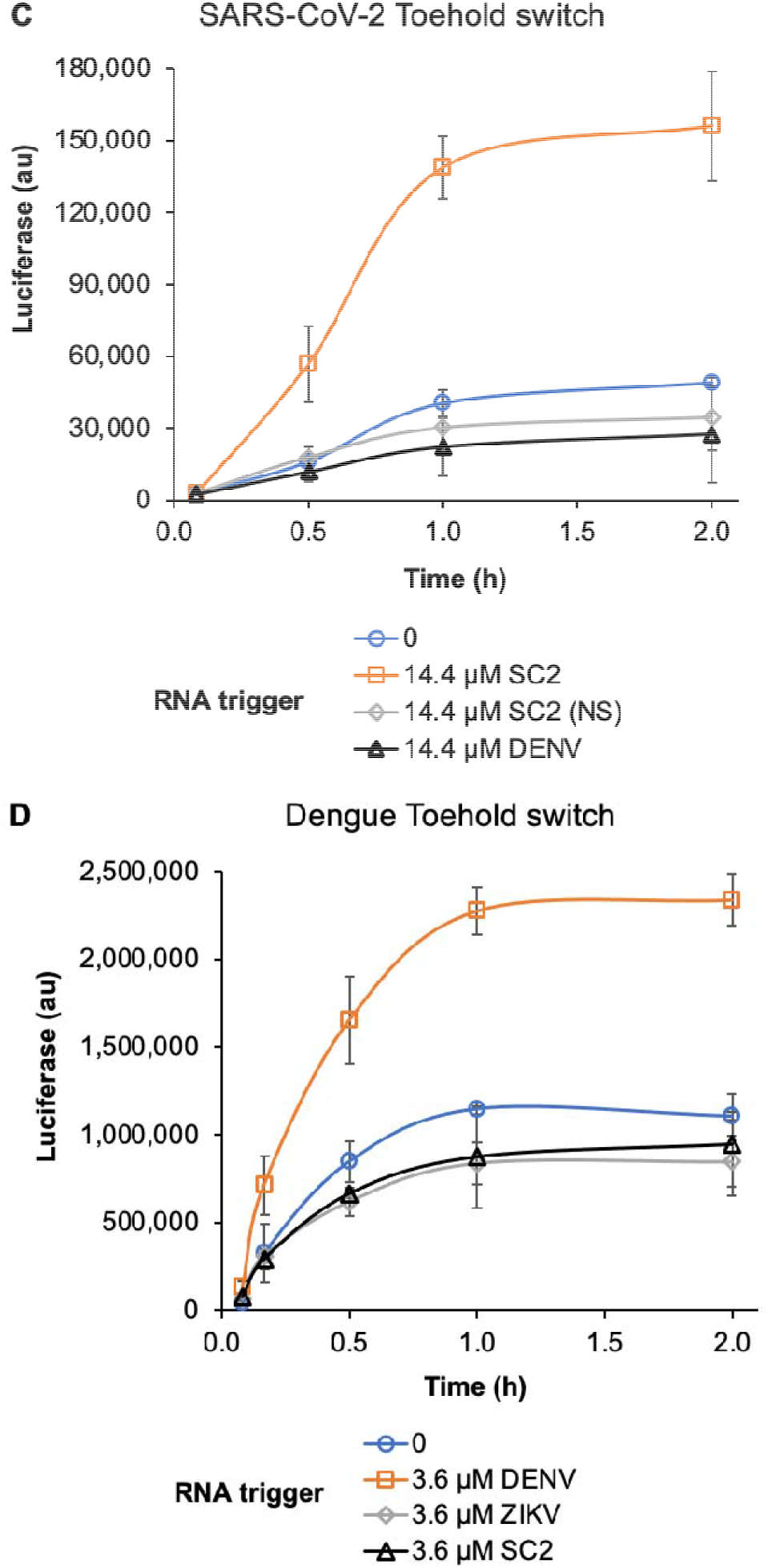
Toehold switches control of the luciferase reporter in cell-free gene expression. (A) The toehold switches prevent the ribosome to access the ribosome binding site (RBS). In the presence of the trigger RNA, base-pairing between the switch and trigger releases the RBS and activates the luciferase gene translation. (B) Switch B triggered by a synthetic trigger B at different concentrations. (C) SARS-CoV-2 switch specificity tested with a specific SARS-CoV-2 trigger (SC2), a nonspecific SARS-CoV-2 trigger (SC2 (NS)), and the nonspecific Dengue trigger (DENV) at 14.4 µM each. (D) Dengue switch specificity tested with the specific Dengue trigger (DENV), and the nonspecific Zika (ZIKV) and SARS-CoV-2 (SC2) triggers at 3.6 µM each. Lower concentrations of triggers did not result in specific activation (Supplementary Figure 1).

### 2.3 T7 RNA polymerase gene circuit

Based on the great specificity but poor sensitivity of our toehold switches we decided to design a signal amplification circuit by placing the T7 RNA polymerase (T7 RNAP) under the control of the Dengue switch and the luciferase gene under the control of the T7 promoter (P_T7_). Therefore, we used the polymerase to transfer the switch signal to the output unit (Figure 2A). The new circuit was functional and well able to discriminate between the Dengue and Zika RNA triggers with a 4-fold change after 30 min (Figure 2B). Indirect control through the T7 RNAP greatly decreased the expression leakage around one order of magnitude and increased the fold change between ON and OFF states compared to the direct control of the output. However, it did not decrease the sensitivity of the sensor below 3.6 µM. Although we were unable to reach our goal fully, the improvements made drove us to the hypothesis that the approach was right but the tool used was nonoptimal. Therefore, we designed a new circuit, replacing the T7 RNAP with a serine recombinase.

**Figure 2.**
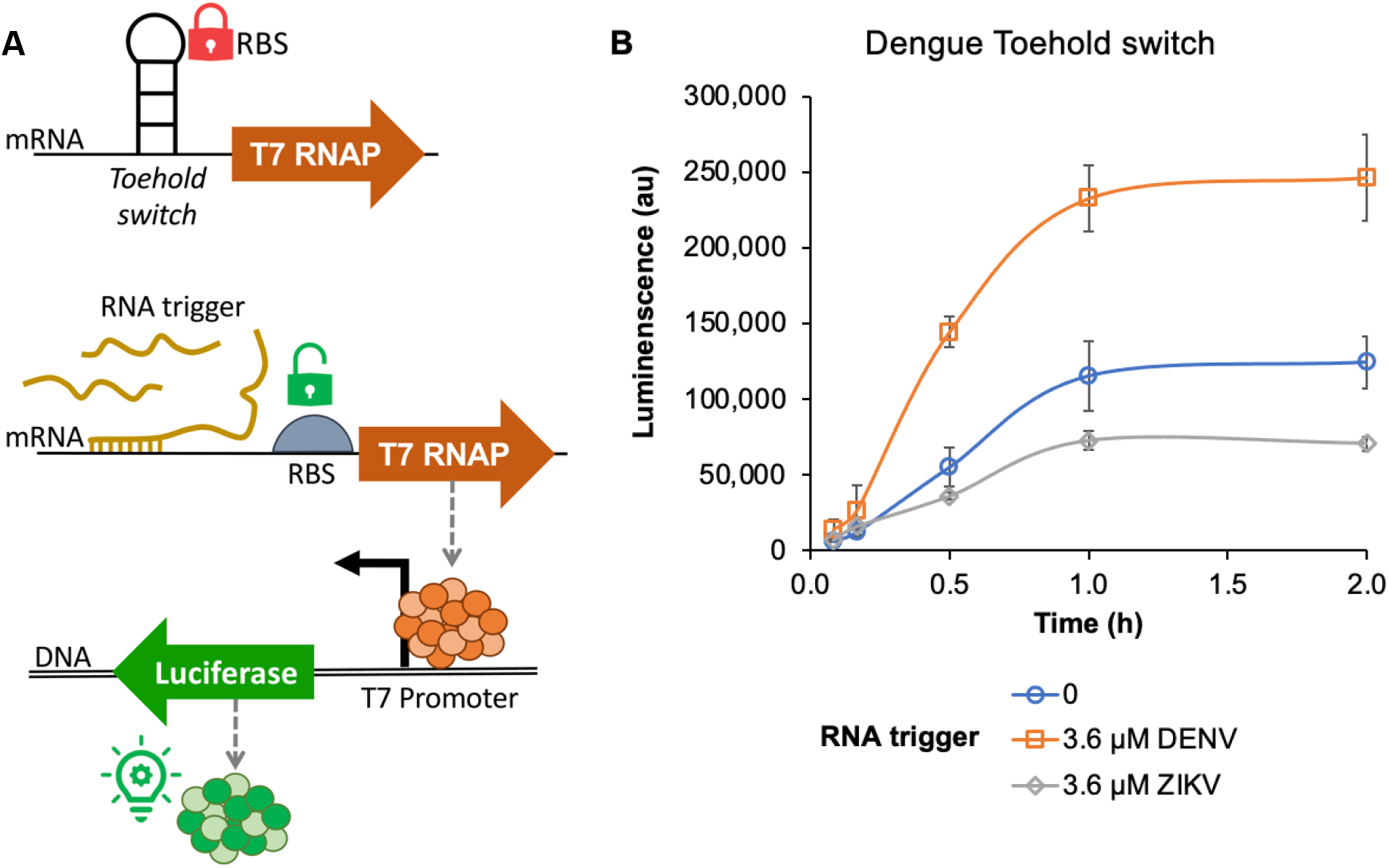
T7 RNA polymerase circuit for signal amplification. (A) The T7 RNAP gene is transcribed from the J23119 promoter and controlled at the translational level by the Toehold switch. Trigger activation leads to the synthesis of the T7 RNAP, which in turn drives the expression of the luciferase gene. (B) Circuit response to the Dengue switch treated with no trigger (0), the specific trigger DENV (Dengue virus RNA), and the nonspecific trigger ZIKV (Zika virus RNA).

### 2.4 Serine-integrase converts the switch response from analogic to digital-like

Serine-integrases are able to flip DNA sequences flanked by the recombination sites *attB* and *attP*. In the new amplification circuit, the integrase 13 ^23,24^ was cloned under the control of the toehold switch in the sensor plasmid, and the luciferase gene was cloned in the response plasmid downstream of the T7 promoter, which was coded in the reverse strand and flanked by the *attB* and *attP* sites. Therefore, switch activation by the trigger should lead to the synthesis of the integrase, which in turn will flip the T7 promoter allowing the expression of the luciferase (Figure 3A). For the convenience of use, we first attempted to clone the complete circuit in one plasmid. However, the switch’s leakage in *E. coli* always resulted in flipped T7 promoter spotted during sequencing for cloning confirmation. Therefore, we divided the circuit into two plasmids, sensor and response. Although less convenient, the two-plasmid device allowed tuning the dose of each plasmid to reach the desired output level. For the more open Dengue switch, using 1:10 sensor to response-plasmid resulted in the best switching activity, while the more closed switches B and Sars-CoV-2 required 1:2 and 1:1, respectively. In the optimized plasmid doses, the new circuits were highly effective in containing leakage (Figure 3B-D), especially the Dengue and the Sars-CoV-2 switches. Moreover, the new circuits present stronger activation of gene expression and higher sensitivity to the RNA trigger. The lowest concentration of trigger detected by all three switches was 7.2 nM, three orders of magnitude lower than the switches’ direct control of the output. Switch control of the integrase delayed the circuit’s response by at least 1h compared to the direct control and the switch-controlled T7RNAP. However, after 2h the circuits strongly activate gene expression showing a digital-like response. The integrase boosted the switches’ dynamic range from up to 3 to 11-21-fold (Figure 3E).

**Figure 3.**
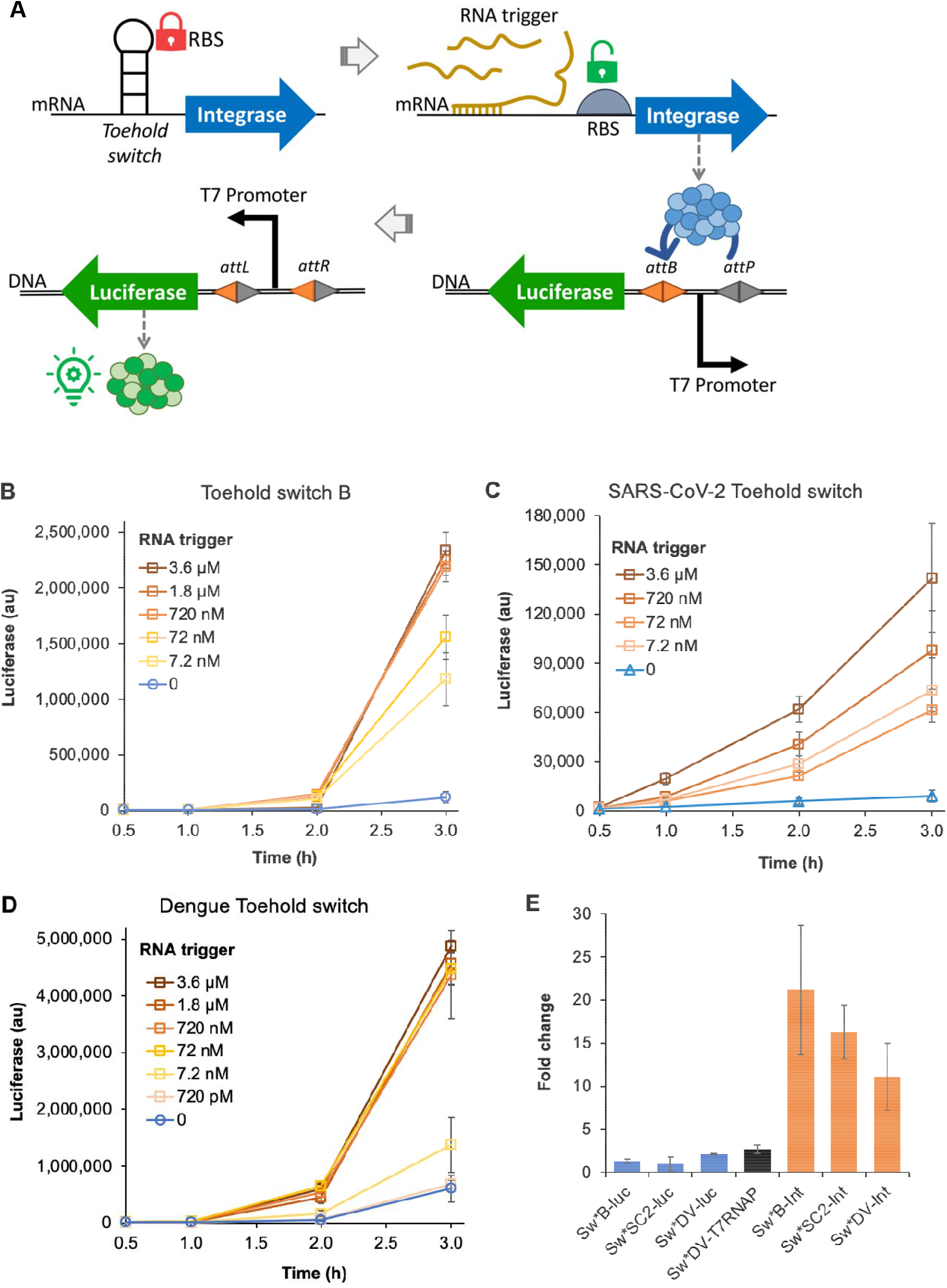
Serine integrase circuit for signal amplification. (A) The integrase gene is transcribed from the T7 promoter and controlled at the translational level by the Toehold switch. The trigger activates the synthesis of the integrase, which flips the T7 promoter in the output DNA unit activating the expression of the luciferase gene. The output unit is composed of the luciferase gene and the T7 promoter coded in opposite senses, which precludes gene expression. (B-D). Circuits response to switches B, SARS-CoV-2, and Dengue treated with no trigger (0), and the specific trigger at different concentrations. (E) Fold-changes for the switches treated with 3.6 µM specific RNA trigger in all constructs tested: switch B (Sw*B), SARS-CoV-2 switch (Sw*SC2), Dengue switch (Sw*DV), luciferase (luc), T7 RNA polymerase (T7RNAP), and serine integrase (Int).

## 3. DISCUSSION

Toehold switches have been used as sensors for the development of cell-free diagnostics after Pardee and collaborators demonstrated that the reaction could be developed on a piece of paper, enabling applications outside lab ^1^. This tool has been developed to respond to important virus outbreaks, such as Ebola ^1^ in West Africa in 2013, Zika ^8^ in the Americas in 2015, and more recently the SARS-CoV-2 pandemic in 2020 ^11,17,25^. A common hurdle for use of toehold switches in cell-free systems is the low dynamic range of the sensors. The best Zika toehold switches activated gene expression up to 30-fold when treat with the cognate trigger; however, from 48 tested sensors 77% did not reach 10-fold change ^8^. For SARS-CoV-2, most switches developed so far show specific activation below 10-fold ^11,17,25^. The highest fold-change of 40 has been demonstrated for one out of 11 switches constructed against N-gene targets from the SARS-CoV-2 genome ^11^. Using sequence-to-function deep learning to design toehold switches, good SARS-CoV-2 sensors initially displayed up to 5-fold activation, and after optimization one reached around 13-fold but most displayed up to 10-fold activation ^25^. Our SARS-CoV-2 switch could reach up to 7.7-fold change only when treated with tens of micromolar concentration of trigger, displaying no activation at a low micromolar concentration.

In a first attempt to amplify the toehold switch’s signal, we tested the T7 RNA polymerase, an enzyme that holds interesting features for the construction of synthetic gene circuits, such as transcriptional activity, orthogonality, and promoter malleability ^26^. The circuit proved functional and able to improve the fold-change by 42%, with significant containment of leakage. However, no sensitivity improvement was achieved. Previous efforts to construct a T7 RNA polymerase cascade circuit controlled by a different toehold switch also succeeded but achieved the same 4-fold activation ^1^. The results with the T7 RNA polymerase indicated we were in the right direction, but needed a more suitable tool for the task. Therefore, we constructed a new circuit using serine integrase. The integrase amplification circuit boosted the switch’s performance 16-fold and improved its sensitivity four orders of magnitude. Such a sensitivity improvement has only been achieved before using a previous step of nucleic acid sequence-based amplification-mediated RNA amplification (NASBA) ^8^. Our amplification circuit is still compatible with any previous sample amplification step, which can potentially boost the system’s sensitivity even further. We demonstrated that the integrase circuit is suitable for different toehold switches. Furthermore, we successfully constructed the first Dengue toehold switch, which is able to discriminate against Zika and SARS-CoV-2 virus RNAs and detects the specific Dengue RNA at low nM concentration when coupled with the integrase amplification circuit.

We developed a simple and efficient circuit to amplify a biosensor’s signal in a cell-free expression system. Improving the performance of genetically encoded sensors can often lead to a trade-off between sensitivity and background expression ^6^. We managed to create a circuit with a very low background expression while displaying an improved output signal and dynamic range. The digital-like response of the integrase circuit is widely useful for RNA and other small molecule sensors, as well as for processing information in synthetic cells. Finally, serine integrase 13 is highly active in bacterial ^24^, mammal, and plant cells ^23^, and in a cell-free system, which makes it a modular and robust tool for engineering circuits.

## 4. MATERIAL & METHODS

### 4.1 Design of toehold switches

The toehold switches were designed and analyzed using the NUPACK Web Application ^15^. A script was developed based on previously published design principles for toehold switches ^12^. Pre-defined 40nt-long sequences from Dengue type 1 and Sars-CoV-2 genomes were used as RNA triggers. Additionally, the script was used to design switch/trigger pairs without trigger sequence constrain. Up to ten switch sequences designed for each target RNA were analyzed. Only switch/trigger pairs showing 100% complex formation in the equilibrium at equimolar concentrations were selected. Next, the switches were analyzed for translation initiation rates in the OFF and ON states using the RBS calculator ^19–21^. Switches predicted to have translation rates up to 10^1^ in the OFF state and equal to or higher than 10^3^ in the ON state were selected and ranked according to their predicted dynamic range (calculated by subtracting the translation rate in the OFF state from the translation rate in the ON state). The best-performing pair switch/trigger for each target was constructed after *in silico* analysis. Trigger and Switch sequences used are presented in Supplementary Tables 1 and 2.

### 4.2 Plasmid and DNA sequences

The plasmid pSB1C3 was used as the backbone of all gene circuits (http://parts.igem.org/Part:pSB1C3). The EPIC Firefly Luciferase gene was PCR amplified from BBa_K325109 and the T7 RNA polymerase from the gDNA of *E. coli* BL21. The forward primers were designed to omit the start codon for both genes. The serine integrase 13 gene was synthesized without the start codon. The PCR products and the synthetic gene were assembled into the pSB1C3 using BioBrick standard cloning. Sequences composed of the T7 promoter and the switches were synthesized and cloned into the plasmids pSB1C3-EPICluc and pSB1C3-Integrase upstream of the reporter genes using *Eco*RI and *Bsa*I restriction sites. Sequences composed of the J32119 promoter and the switch were also synthesized and cloned into the pSB1C3-T7RNAP upstream of the reporter gene using *Eco*RI and *Bsa*I restriction sites.

The EPIC Firefly Luciferase was PCR amplified again, this time including the start codon, using a modifying forward primer to insert the T7 promoter and an RBS and assembled into the pSB1C3-T7RNAP using BioBrick standard cloning to construct the T7 RNA polymerase circuit plasmid pSB1C3-T7RNAP-EPICluc. The luciferase gene was PCR amplified a third time using a modifying forward primer to the RBS only. The resulting PCR product and a synthetic sequence encompassing the *attP* site, the T7 promoter coded in the reverse strand, the T500 double terminator, and the *attB* recombination site were assembled into the pSB1C3 using BioBrick standard cloning to construct the integrase response plasmid. All the plasmids used are described in Supplementary table 3.

*E. coli* Top10 was used for cloning and propagation of all plasmids. It was aerobically cultivated at 37°C in Lysogeny Broth (LB) enriched with 25 µg/mL chloramphenicol.

### 4.3 Synthesis of RNA triggers

Oligonucleotides forward and reverse corresponding to the T7 promoter-controlled trigger sequences were annealed and used as templates for *in vitro* transcription. Reactions were set using the RiboMAX™ Large Scale RNA Production System (Promega) and incubated at 37ºC overnight. Reaction inactivation was performed at 70°C for 20 min. Samples were subsequently treated with RQ1 RNase-Free DNase (Promega) for 2h at 37ºC. Finally, trigger RNAs were purified through ethanol precipitation and resuspended in RNase-free water.

### 4.4 *In vitro* gene expression

Template plasmids were extracted from overnight cultures of *E. coli* by miniprep (PureYield™ Plasmid Miniprep System, Promega), ethanol precipitated, and resuspended in RNase-free water. *In vitro* gene expression was carried out using either the *E. coli* T7 S30 Extract System for Circular DNA or the *E. coli* S30 Extract System for Circular DNA (Promega). The second kit was only used for the T7 RNA polymerase circuit. Reactions were set up by adding 1 μL of 0.25 mM amino acid mix, 4 μL of S30 Premix, 3 μL of T7 Extract (or S30 extract), 1 μL of RNA trigger solution, and 1 μL of template plasmid. Plasmids were used at the following concentrations: 2 ng/μL Switch B-luciferase, 2 ng/μL SARS-CoV-2 switch-luciferase, 10 ng/μL Dengue switch-luciferase, 10 ng/μL Dengue switch-T7 RNAP-luciferase, 5 ng/μL Switch B-integrase, 5 ng/μL SARS-CoV-2 switch-integrase, 1 ng/μL Dengue switch-integrase, and 10 ng/μL luciferase circuit response plasmid. The reaction mixture was incubated at 30 °C up to 3h. At defined time points 1 μL samples were withdrawn and mixed with 9 μL of 1% (m/v) bovine serum albumin to stop the reaction.

### 4.5 Luminescence measurements

Luciferase activity was assayed using samples from *in vitro* gene expression reactions. 10 μL of stopped reaction samples were mixed with 50 μL of luciferase substrate solution (Luciferase Assay System kit, Promega) in a white flat bottom 96-well microplate. Luminescence emission was measured in a microplate reader (Tecan Infinite 200 Pro, Switzerland) at 26°C, set for 1000 ms integration time.

## Author Contribution

R.A.L.F.: conceptualization, investigation, data curation, and analysis. G.B.: investigation, data curation, and analysis. V.F.B.Z.: investigation and validation. G.B.P.: investigation and validation.

R.N.L.: conceptualization. E.L.R.: funding acquisition, supervision, and writing - review & editing.

D.B.P.: conceptualization, funding acquisition, project administration, writing - review & editing.

M.R.C.R.L.: conceptualization, supervision, investigation, and writing - original draft.

## Supporting Information

Supplementary material: nucleic acid sequences and plasmids used in this work, supplementary results, and NUPACK script (DOC).

Supplementary data: Data 1 - Raw and processed data for toehold switches’ direct control of the luciferase gene; Data 2 - Raw and processed data for the T7 RNA polymerase circuit; Data 3 - Raw and processed data for the serine integrase circuit (Excel files).

## Acknowledgment

This work was supported by the São Paulo Research Foundation (FAPESP) [grants 2016/02516-2, 2016/02520-0, 2018/24272-3]; Conselho Nacional de Desenvolvimento Científico e Tecnológico (CNPq) [grants 401990/2016-8, 310023/2020-3, and 465603/2014-9]; and Coordenação de Aperfeiçoamento de Pessoal de Nível Superior - Brasil (CAPES) [grant 88887.468198/2019-00, 88887.596758/2020-00, and Finance Code 001]. E.L.R. is supported by Embrapa Genetic Resources and Biotechnology/National Institute of Science and Technology in Synthetic Biology, National Council for Scientific and Technological Development (465603/2014-9), Research Support Foundation of the Federal District (0193.001.262/2017).

