## Supplementary material for "Signal-amplification for cell-free biosensors, an analog-to-digital converter"

*Contributed equally as first author

^†^Corresponding authors:

**Supplementary Table 1**. Sequences corresponding to RNA triggers used in this study

| **Trigger name** | **Origin** | **Sequence** |
| --- | --- | --- |
| B | NUPACK generated | ACUCCUACCAAUCAUCCUACCUACCUUACUUCACGCCCUC |
| DENV | Dengue type 1 virus genome (GenBank: KP188567.1) | AAGCUGUACGCAUGGGGUAGCAGACUAGCGGUUAGAGGA |
| ZIKV | Zika virus genome (GenBank: OK571913.1) | UUCUAGAGAUGCAAGACUUGUGGCUGCUGCGGAGGUCAGA |
| SC2 specific | SARS-CoV-2 virus genome (GenBank: MT126808.1) | AUUUACAACCAUUAGAACAACCUACUAGUGAAGCUGUUGAA |
| SC2 nonspecific | SARS-CoV-2 virus genome (GenBank: MT126808.1) | AUUGUACAGAAAGUGUGUUAAAUCCAGAGAAGAAACUGGCC |

**Supplementary Table 2**. Sequences corresponding to RNA switches used in this study

| **Switch name** | **Sequence** |
| --- | --- |
| Switch B | GAGGGCGUGAAGUAAGGUAGGUAGGAUGAUUGGUAGGAGUGAGUAACGAAAGCCAGAGGAGUUACUAUGUCCUACCAAUCAUCCUAC |
| Dengue Switch | UCCUCUAACCGCUAGUCUGCUACCCCAUGCGUACAGCUUGUUAUAGUUAUGAACAGAGGAGACAUAAUAUGACUAAGCUGUACGCAGUC |
| SARS-CoV-2 Switch | UUCAACAGCUUCACUAGUAGGUUGUUCUAAUGGUUGUAAAGUGUAUAACAGGAGGAAUACAAUGUACAACCAUAAGAACAAC |

**Supplementary table 3**. Plasmids used in this study

| **Plasmid name** | **Description** | **Ref** |
| --- | --- | --- |
| pSB1C3-mRFP1 | Backbone plasmid | BBa_J04450 |
| pSB1C3-EPICluc | EPIC Firefly Luciferase gene template | BBa_K325109 |
| pSB1C3-PT7-EPICluc | EPIC Firefly Luciferase gene under the control of the T7 promoter and an RBS | This work |
| pSB1C3-SwitchB-EPICluc | EPIC Firefly Luciferase gene under the control of the T7 promoter and the Switch B | This work |
| pSB1C3-DENVSwitch-EPICluc | EPIC Firefly Luciferase gene under the control of the T7 promoter and the Dengue switch | This work |
| pSB1C3-SARS-CoV-2Switch-EPICluc | EPIC Firefly Luciferase gene under the control of the T7 promoter and the SARS-CoV-2 switch | This work |
| pSB1C3- DENVSwitch-T7RNAP | T7 RNA polymerase gene under the control of the J32119 and the Dengue switch | This work |
| pSB1C3- DENVSwitch-T7RNAP-EPICluc | T7 RNA polymerase gene under the control of the J32119 and the Dengue switch, and the EPIC Firefly Luciferase gene under the control of the T7 promoter | This work |
| pSB1C3- SwitchB-Integrase | Serine integrase 13 gene under the control of the T7 promoter and the Switch B | This work |
| pSB1C3- DENVSwitch -Integrase | Serine integrase 13 gene under the control of the T7 promoter and the Dengue switch | This work |
| pSB1C3- SARS-CoV-2Switch -Integrase | Serine integrase 13 gene under the control of the T7 promoter and the SARS-CoV-2 switch | This work |
| pSB1C3-attP<PT7(rev)-T500>attB-EPICluc | EPIC Firefly Luciferase gene under the control of the T7 promoter, which is encoded in the reverse strand together with the T500 terminator and is surrounded by the recombination sites *attP* and *attB* (integrase response plasmid) | This work |

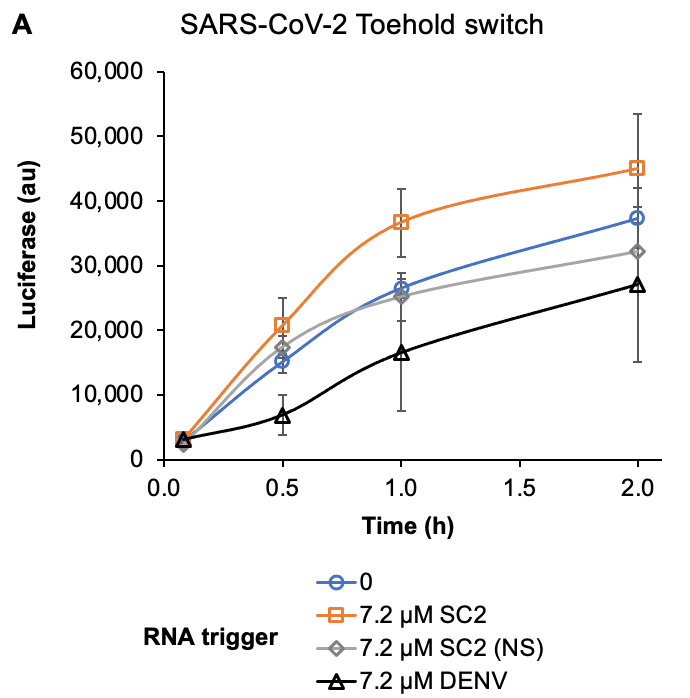

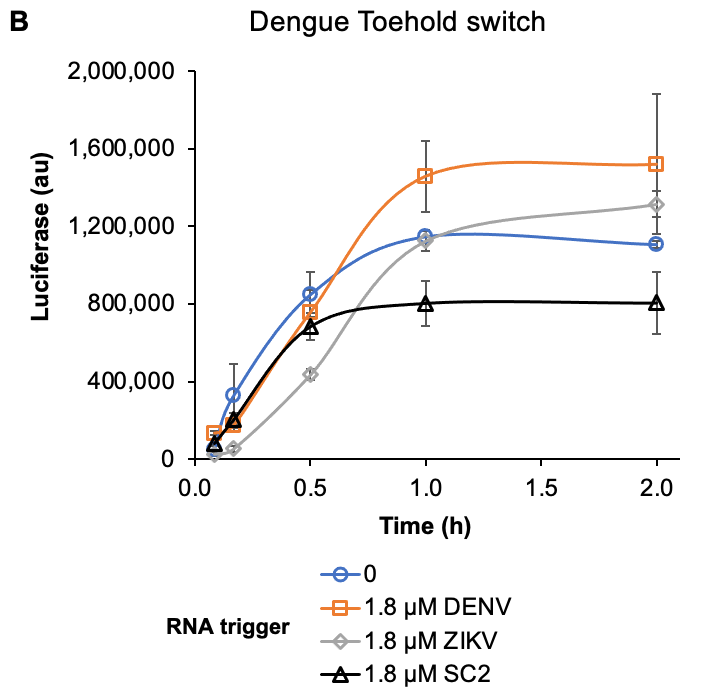

**Supplementary Figure 1**. Toehold switches control of the luciferase reporter in cell-free gene expression. (A) SARS-CoV-2 switch specificity tested with a specific SARS-CoV-2 trigger (SC2), a nonspecific SARS-CoV-2 trigger (SC2 (NS)), and the nonspecific Dengue trigger (DENV) at 7.2 µM each. (D) Dengue switch specificity tested with the specific Dengue trigger (DENV), and the nonspecific Zika (ZIKV) and SARS-CoV-2 (SC2) triggers at 1.8 µM each.

**NUPACK script**

material = rna

temperature = 30

\# target structures

structure switch = U20 D18 (U3 D5 U15 U3)

structure trigger = U40

structure detect = D40 + U47

\# sequence domains

\# a* = trigger

\# if trigger-constrained, past reverse complement in the domain a (N40)

\# domain c = RBS + initial codon

domain a = N40

domain b = N6

domain c = N3 VVVVVAGAGGAGWWW N2 AUG

domain d = N18

\# thread sequence domains onto target structures

switch.seq = a b c d

trigger.seq = a*

detect.seq = a* a b c d

\# specify stop conditions for normalized ensemble defect

\# default: 1.0 (percent) for each target structure

trigger.stop = 5.0 # larger defect for unpaired structures

prevent = AAAA, CCCC, GGGG, UUUU, KKKKKK, MMMMMM, RRRRRR, SSSSSS, WWWWWW, YYYYYY
